# *In-silico* analysis reveals hub genes and enriched pathways in Type 2 Diabetes Mellitus

**DOI:** 10.1101/2023.08.21.553951

**Authors:** Prasad Gandham, RG Sharathchandra

## Abstract

The worldwide prevalence of diabetes mellitus poses a significant hazard to human health and a substantial economic burden for modern society. Multiple genes and gene-environment interactions make up the complicated genetic landscape of type 2 diabetes (T2D). The majority of genetic studies concentrated on separate genes, while some put an excessive amount of emphasis on SNPs. The objective of this work was to analyze all known T2D-associated genes in a collective manner in order to identify candidate genes and assess their possible functional role using bioinformatic tools. Interactome networks, enrichment clustering, hub genes, enriched biological pathways, and miRNAs targeting hub genes were analyzed. From the pool of 490 genes implicated in type 2 diabetes, the 25 most prominent hub genes were identified using measures of betweenness centrality and degree of each node. From the interactome analysis 8 different MCODE clusters was observed with a highest number of 63 genes in cluster one. GO analyses revealed that candidate genes were significantly enriched in glucose homeostasis, carbohydrate homeostasis, regulation of type B pancreatic cell apoptotic process, and regulation of hormone secretion biological process. Insulin resistance, PI3K-Akt signaling pathway, AMPK signaling pathway, Neuroactive ligand-receptor interaction, and Adipocytokine signaling pathway were among the notable enriched pathways. Subsequently we found 54 miRNAs targeting 25 hub genes out of which IRS1 and IGF1 hub genes were targeted by seven different miRNA. This study contributes to a thorough bioinformatics investigation of genes, functions, and pathways that are relevant to the etiology of T2DM and may be useful in the development of precision medicine-based treatments.

## Introduction

Diabetes mellitus (DM) is a chronic metabolic condition that occurs as a result of hyperglycemia (high blood glucose levels)[1], happen when the pancreas is no longer able to produce enough of the hormone insulin, or when the body cannot effectively use the insulin it produces. According to recent data from the International Diabetes Federation (IDF) diabetes atlas tenth edition 2021, there were 537 million diabetic adult (20-79 years) patients worldwide and this number is expected to rise to 783 million by 2045. It has been reported that youth from Canadian first nations, American Indian, Australian aboriginal and African american populations are having highest incidence rate of T2DM[2] [3] [4] [5]. Whereas non-Hispanic Caucasian youth populations, such as those in the Europe and US had the lowest incidence rates [6] [7]. A 2019 study by the Centers for Disease Control and Prevention (CDC) revealed that the prevalence of type 2 diabetes among racial/ethnic groups in the United States was as follows White population 12.1%, Asian population 19.1%, African Americans 20.4%, and Latinx people 22.1% [8]. It is important to note that these statistics are constantly changing as new data becomes available, and the prevalence and impact of Type 2 Diabetes can vary greatly depending on the country and population being considered. Type 1 diabetes, type 2 diabetes, and pregnancy induced diabetes are the main types of diabetes in clinical practice, of which around 90 percent are type 2 diabetes mellitus (T2DM) cases [9] [10]. The primary risk factors for Type 2 Diabetes include obesity, high blood pressure, physical inactivity, unhealthy diet, and increasing age [11]. Besides hyperglycemia, T2DM related potential complications include nephropathy, neuropathy, retinopathy, and cardiovascular disease, making it the leading causes of death worldwide [12] [13] [14].

Although Type 2 diabetes is a complex metabolic condition that is influenced by ethnicity, age, behavioral and environmental factors, genetic susceptibility plays a larger role than others due to the incidence of high-risk genes. Which is backed up by the fact that in large cohort of family and twin studies showed that heritability of type 2 diabetes estimated to be in the range of 25% to 72% [15]. Recent data suggests that both insulin secretion and resistance are inherited features, and studies have established a genetic connection to faulty insulin secretion and resistance in type 2 diabetics [16] [17]. Different Genome-wide association studies (GWAS) have identified multiple susceptibility loci for type 2 diabetes and related traits like insulin resistance and beta-cell dysfunction [18] [19]. However, because imputation of ungenotyped variants was dependent on accessible reference panels from resources like as the HapMap, many of these GWAS only identified common variants. The mechanisms by which these susceptibility loci affect and cause T2DM remain unclear.

The aim of this study is to perform a collective computational analysis of T2DM biomarker panel consisting of all documented associated genes and their corresponding proteins using enrichment and interaction tools to prioritize candidates. We established a protein–protein interaction (PPI) network of the T2DM biomarker panel, and hub genes were showcased. Under enrichment analysis we performed functional annotation analysis using the Gene Ontology (GO) and pathways enrichment analysis using the Kyoto Encyclopedia of Genes and Genomes (KEGG), used to determine underlying biological processes and biochemical pathways relative to the hub genes. The present study may be useful to explore potential targets and concurrent designing of novel strategies for clinical therapies. The insulin resistance a pathophysiological mechanism contributing to T2DM, has a significant genetic component [20].

## Materials and methods

### Identification of T2D-associated genes and preparation of datasets

The public free scientific and genomic repositories like PubMed (Lindberg 2000) and NCBI (http://ncbi.nlm.nih.gov) were used to retrieve T2DM-associated genes from human studies. Studies linked to T2D were searched in PubMed using the terms ‘Type 2 Diabetes & Genotype’, ‘Type 2 Diabetes & Alleles’ and ‘Type 2 Diabetes & Polymorphisms genetic’. A total of 5281, 2810 and 4291 respective publications were retrieved from PubMed. There was an overlap of 3941 publications. All the search items from PubMed were thoroughly sorted, and research reporting gene(s) associated with T2D regarding discovery, identification, genetics, GWAS, polymorphisms, clinical, biochemical, molecular and pathophysiological conditions etc. were incorporated so as to ensure data reliability. Finally, a total of 490 genes were retrieved and curated into a dataset, viz. T2D-associated genes (Supplementary Taable 1).

### Interactome analysis

In order to study the protein-protein interactions between T2D genes we have used Metascape framework, this intern contains data from STRING [21], BioGrid [22], InWeb_IM [23] and OmniPath [24] interactome databases. The effective data driven gene prioritization was performed using Metascape [25] from set of gene and protein identifiers, extracted rich annotations, and constructed PPI networks. By combining data from above mentioned databases Metascape produced four datasets namely Physical (Core), Physical (All), Combined (Core), and Combined (All). Currently Metascape contains more than 2.8 million human physical PPI pairs in which >2 million were from STRING database. Integration of data from four different resources into Metascape enabled room for quick and extensive analysis. The robust and computationally expensive algorithms diploid in Metascape predicts protein interaction networks. In order to obtain more biologically interpretable results Metascape employs MCODE [26], a mature complex identification algorithm to recognize neighbourhoods or closely connected protein complexes embedded in large protein networks. The Cytoscape v3.7.1 [27] was used to visualize the MCODE generated protein networks with forced directed layout. The all the genes included in this study and their interactions were visualized as a Circos [28] plot.

### Functional enrichment analysis

To identify enriched GO terms in a list of all candidate genes compared to a background list of genes Gene ontology enrichment analysis and visualization tool (Gorilla) [29] was used. The Kyoto Encyclopedia of Genes and Genomes (KEGG) Orthology-Based Annotation System (KOBAS-i) [30] was used to calculate the enriched KEGG pathways from candidate genes. The statistical test used for enrichment analysis is typically based on a hypergeometric test and for false discovery rate (FDR) correction Benjamini-Hochberg [31] method was applied. The terms with p-value less than 0.01, FDR cutoff of 0.01 and a minimum count of two gene involved were considered significantly enriched among the test set. These thresholds were used to avoid overlap and minimize redundancy in ontology terms. All gene identifiers in the genome have been used as the enrichment background reference set. The top 25 statistically significant phrases with the best p-value were chosen as representatives and depicted in the GO and KEGG enrichment figures.

### Identification of miRNAs targeting hub genes

The microRNAs (miRNAs) targeting the predicted hub genes were identified using the psRNAtarget web-based tool [32]. It predicts the target genes based on complementarity matching between the miRNA and the target mRNA sequence using predefined scoring schema and evaluating target site accessibility. To get high confidence miRNA target genes maximum expectation value was set to 3. A smaller expectation value indicates high complementarity between miRNA and target gene. Hub genes and miRNA interactions were plotted as a Gene-miRNA interaction network using Cytoscape software.

### Results

We collected a total 490 associated genes from different studies on T2D, as shown in the figure 1 these genes were found to be distributed on 1–22 (chr1–chr22) autosomes, one sex chromosome X (chrX) as well as on the Mitochondrial (M) genome. The most abundant genes are located on chromosome 1, containing 49 candidate genes, followed by chromosomes 11 and 2 containing 44 and 36 candidate genes respectively. Additionally, 35, 31 and 31 number of candidate genes were found to be located on chromosomes 6, 7 and 12 respectively. The 11 genes are also found to be located on chromosome X and 2 are on mitochondrial genome. From the distribution of multiple genes on different chromosomes it is clearly evident that T2D is a complex and polygenic disease. The ideogram of Circos plot representing genomic location of all the genes included in this study and their interactions were shown as links in the central portion of the plot (Figure 2). The color of each link corresponds to the MCODE subnetwork color.

**Figure 1:**
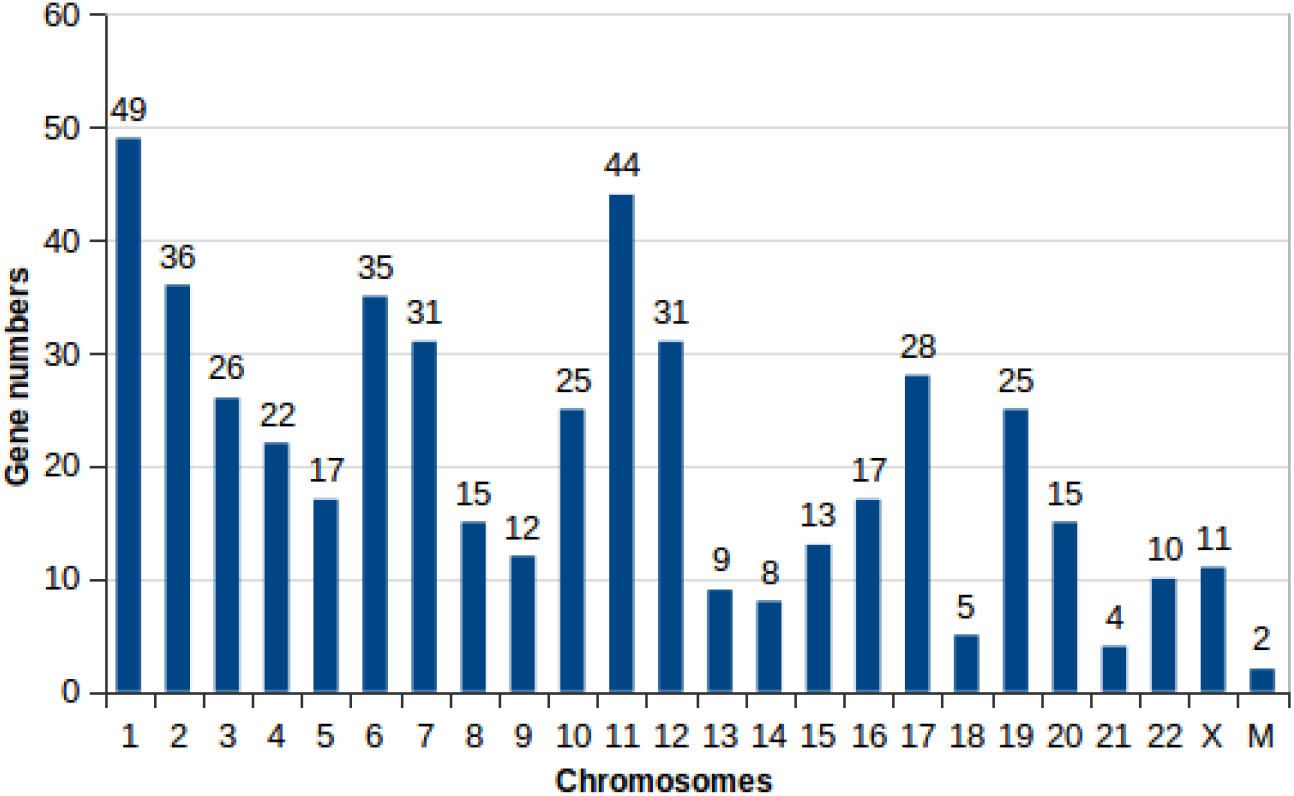
Distribution of 490 genes included in this study were shown on human chromosomes

**Figure 2:**
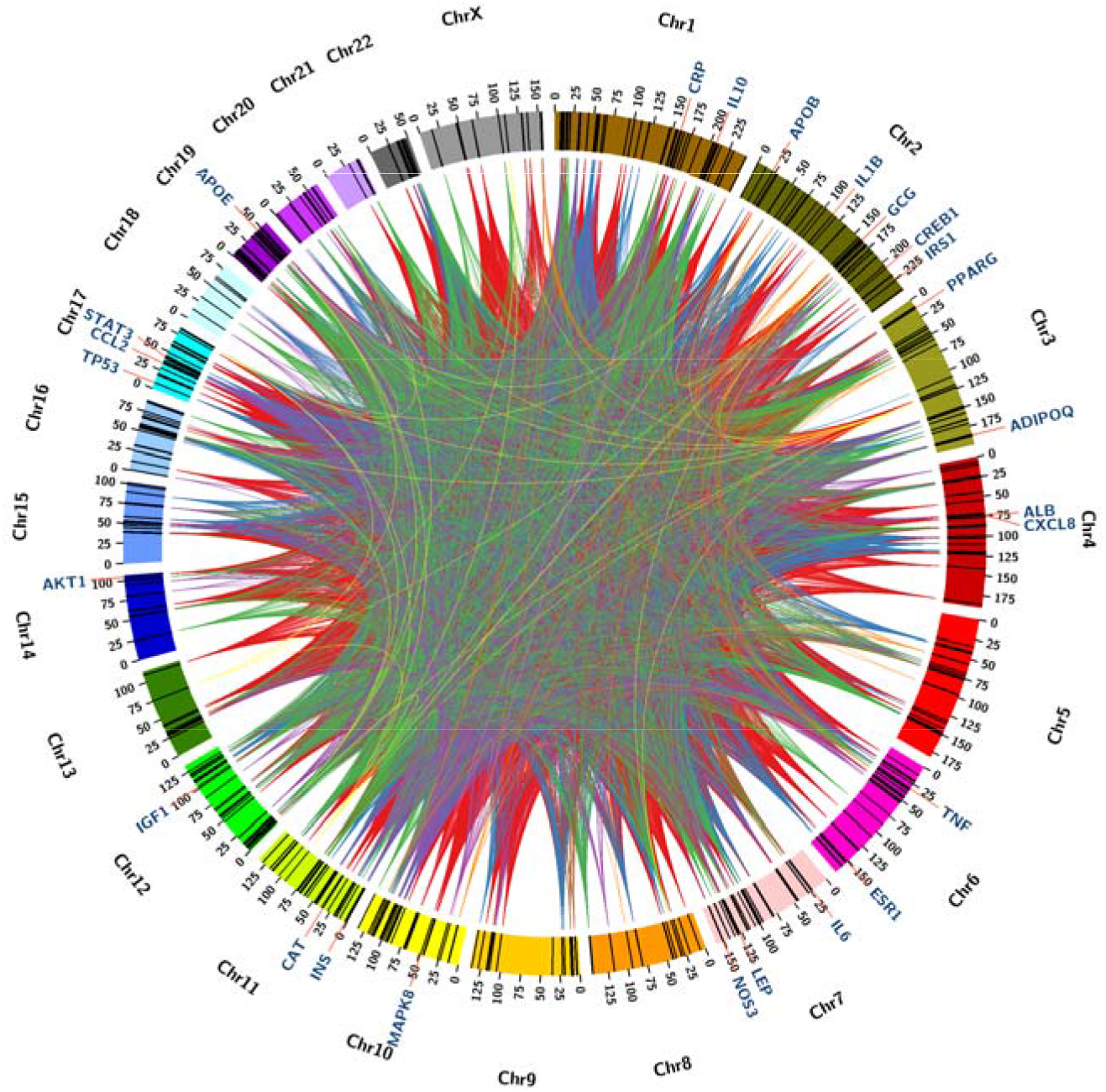
Circos plot depicting genome features across the 23 human chromosomes.

### PPI network of T2D associated genes and analysis of modules

The protein-protein interactions (PPI) network analysis of the T2D associated genes may provide insight into key biochemical systems controlling biological processes. The PPI networks are graphical representations of the interactions between proteins within a cellular system. The 490 candidate genes in T2D were subjected to Metascape PPI network analysis and the resulted massy hairball network consists of 4036 number of edges. To get more biologically interpretable outcomes subnetworks embedded in the large network were extracted by applying MCODE algorithm. This algorithm was used to precisely detect and categorize densely connected regions containing molecular complexes present in protein-protein interactions. The eight detected MCODE components/clusters from T2D associated genes were portrayed with unique color in figure 3. Each node is labeled with its respective gene name and its size correlated with statistical significance and hubs of network. There were 196 genes in total among the 8 MCODE components. The number of genes found in Clusters 1 through 8 are as follows 63, 45, 24, 19, 18, 14, 9, and 4, respectively. Clusters 1 and 2 had the highest number of genes with 63 and 45 genes, 553 and 201 interactions (Edges) respectively, while MCODE clusters 7 and 8 had the lowest share of 9 and 4 genes, 35 and 5 interactions. The genes in different MCODE clusters were illustrated in supplementary table 2.

**Figure 3:**
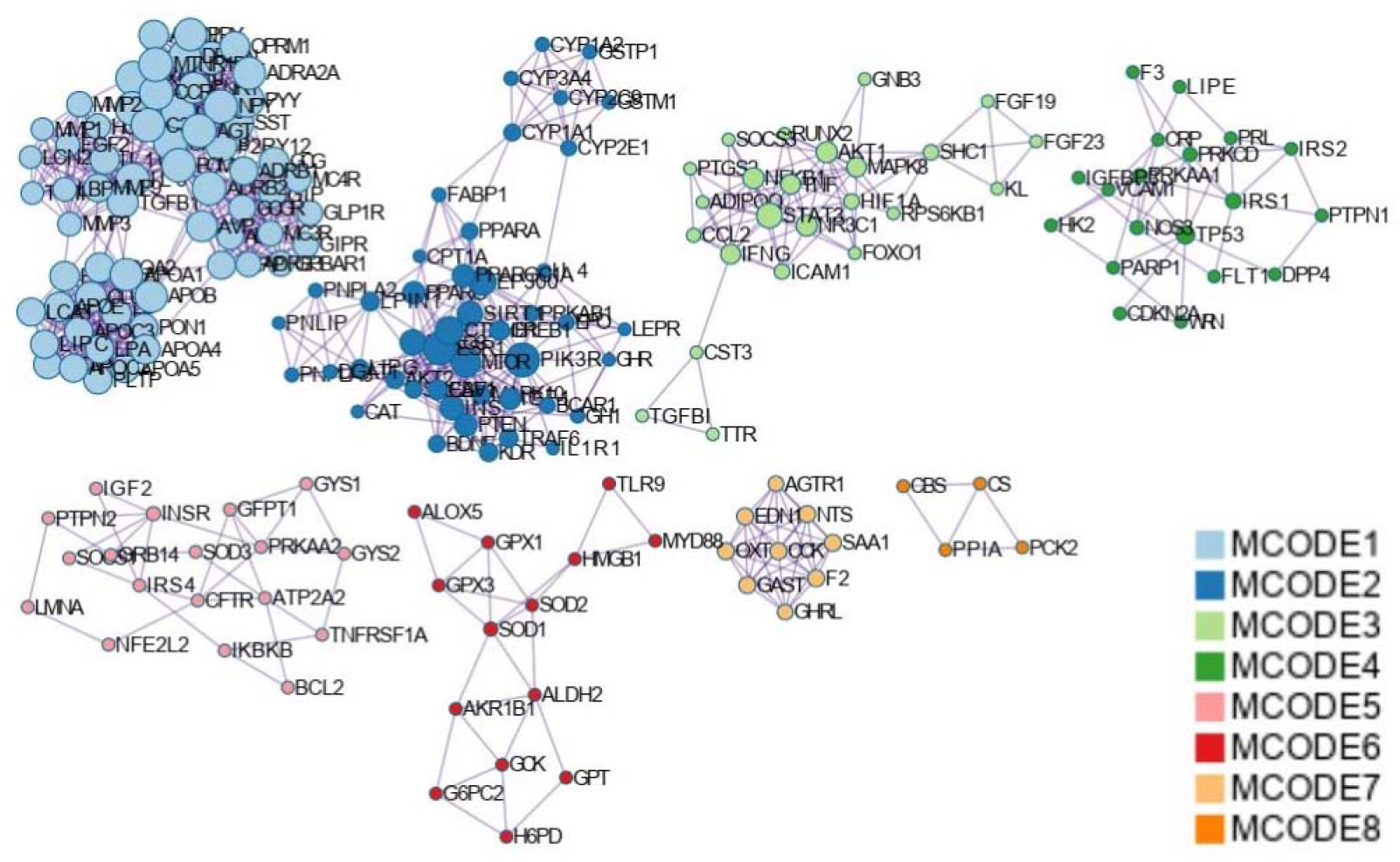
The MCODE subnetworks from the list of T2D-associated genes and each subnetwork had a different unique color. Each node is labeled with gene symbol and node size correlated with statistical significance and hubs of network. The 8 MCODE components contained 429 genes. https://metascape.org/gp/index.html#/reportfinal/tssqhqy41

Each MCODE component was analyzed independently using pathway and process enrichment analysis, and the top three enriched terms by p-value were listed and functional description of the corresponding components given in the supplementary table 3. It was found that the most significantly enriched terms in component one consists of GPCR ligand binding, Signaling by GPCR and GPCR downstream signaling. G protein-coupled receptors (GPCRs) are involved in a wide variety of physiological functions, including glucose homeostasis and insulin production [33]. The beta-2 adrenergic receptor (β2AR) is one of the most studied GPCRs in type 2 diabetes, β2AR gene knockout mice showed a reduced glucose-stimulated insulin release by pancreatic β-cells that leads to deterioration of glucose tolerance [34]. The genes of the MCODE component 2 were mainly involved in AMPK signaling pathway, Alcoholic liver disease and Longevity regulating pathway. AMP-activated protein kinase (AMPK) activation evoke insulin-sensitizing effects, when cellular energy levels are low, and it signals to stimulate fatty acid oxidation in adipose tissues, and reduces hepatic glucose production. Brain-derived neurotrophic factor (BDNF) plays a crucial role in longevity regulation by protecting against age related cognitive decline and neurological disorders, as well as its potential to promote cellular and mitochondrial[35] health [36] [37]. The regulation of small molecule metabolic process, Signaling by Interleukins, Interleukin-4 and Interleukin-13 signaling are the major enriched pathways in component three. The FGF19 (fibroblast growth factor 19) and FGF23 (fibroblast growth factor 23) helps keep the balance of glycolipids and may involve in regulating the secretory activity of islet beta cells in patients with type 2 diabetes [38]. ATP2A2, BCL2, CDKN2A, CFTR, CRP, DPP4, F3, FLT1, GFPT1, GRB14, GYS1, GYS2, HK2, IGF2, IGFBP3, IKBKB, INSR, IRS1, IRS2, IRS4 etc., revealed major pathways like Insulin resistance, Insulin signaling pathway, and Focal adhesion: PI3K-Akt-mTOR-signaling pathway in the MCODE components 4 and 5 IRS1 (Insulin Receptor Substrate 1) is signal transduction proteins activated in response to insulin binding to its receptor on the cell surface, and play a crucial role as signaling intermediate which facilitate the translocation of glucose transporters to the cell membrane, allowing glucose to enter the cell [39]. IRS2 is essential for beta-cell under conditions of elevated insulin need, beta-cell failure in human T2D likely results from altered IRS2-dependent signaling, and islets extracted from T2D patients show lower IRS2 expression [40]. MCODE component 6 revealed the importance of selenium micronutrient network, oxidative stress response, and detoxification of reactive oxygen species (ROS) enriched by GCK, H6PD, AKR1B1, GPT, GPX3, ALDH2, TLR9, GPX1, SOD2, ALOX5, MYD88, HMGB1, SOD1, and G6PC2 genes. The generation of reactive oxygen species as a result of oxidative stress which occurs when free radicals are generated in the body has been proposed as the underlying cause of insulin resistance, beta-cell malfunction, impaired glucose tolerance, and type 2 diabetes [41]. Enrichment of G alpha (q) signaling events, and Peptide ligand-binding receptors in MCODE component 7 was reinforced by CCK, SAA1, AGTR1, OXT, F2, GHRL, NTS, GAST, and EDN1genes. MCODE component 8 housed CS, PPIA, CBS, and PCK2 genes.

### Hub genes identification

To prioritize the candidates the Metascape PPI massy hairball network topological characteristics like Degree (D), BetweennessCentrality (BC), ClosenessCentrality (CC) and AverageShortestPathLength (ASPL) for each candidate gene was estimated using Cytoscape’s Network Analyzer program. The degree centrality and betweenness centrality are two different important measures of network centrality methods used to identify hub genes. Degree centrality is a measure of the number of connections that a node has in the network and Betweenness centrality is a measure of the extent to which a node lies on the shortest paths between other nodes in the network. The distribution of degree and betweenness centrality of each node was plotted as a scatter plot with a regression line as shown in the figure 4. The top five percent of the nodes with high degree and betweenness centrality were highlighted with blue color and those 25 are considered as hub genes. The 25 hub genes namely insulin (INS), AKT Serine/Threonine kinase 1 (AKT1), albumin (ALB), interleukin 6 (IL6), tumor necrosis factor (TNF), peroxisome proliferator activated receptor gamma (PPARG), leptin (LEP), insulin like growth factor 1 (IGF1), adiponectin C1Q and collagen domain containing (ADIPOQ), tumor protein P53 (TP53), apolipoprotein E (APOE), mitogen-activated protein kinase 8 (MAPK8), C-C motif chemokine ligand 2 (CCL2), estrogen receptor 1 (ESR1), insulin receptor substrate 1 (IRS1), glucagon (GCG), signal Transducer and activator of transcription 3 (STAT3), catalase (CAT), interleukin 10 (IL10), C-reactive protein (CRP), CAMP responsive element binding protein 1 (CREB1), nitric oxide synthase 3 (NOS3), C-X-C motif chemokine ligand 8 (CXCL8), apolipoprotein B (APOB), and interleukin 1 beta (IL1B), Table 1 shows their positions on the genome, related transcription factors, degree, and betweenness centrality of each hub gene. The 25 hub genes identified in this study were labeled with their gene symbol in blue color in circos plot. Out of 25 hub genes 5 are on chromosome 2 and 3 each on chromosomes 7 & 17 were located, all of their positions were given in the supplementary table 2. The IL1B, IL6 and IL10 identified as hub genes belons to the signaling molecules interleukins (ILs) that play important role in the immune system and inflammation. In type 2 diabetes, interleukins are involved in the pathogenesis of the disease and its associated complications. The interleukin-6 has been extensively studied in type 2 diabetes, is produced by various cells in the body, including adipose tissue, and it has inflammatory effects. In type 2 diabetes, elevated levels of IL-6 have been observed, and this cytokine has been implicated in the development of insulin resistance that characterizes the disease [42]. On the other hand, Interleukin-10 (IL-10) is an anti-inflammatory cytokine that has a protective role in type 2 diabetes by downregulating the production of key pro-inflammatory cytokines like IL-1, IL-6 and TNF-α. The IL-10 polymorphism (rs1800896) appears to play a major role in the development of type 2 diabetes [43].

**Table 1:**
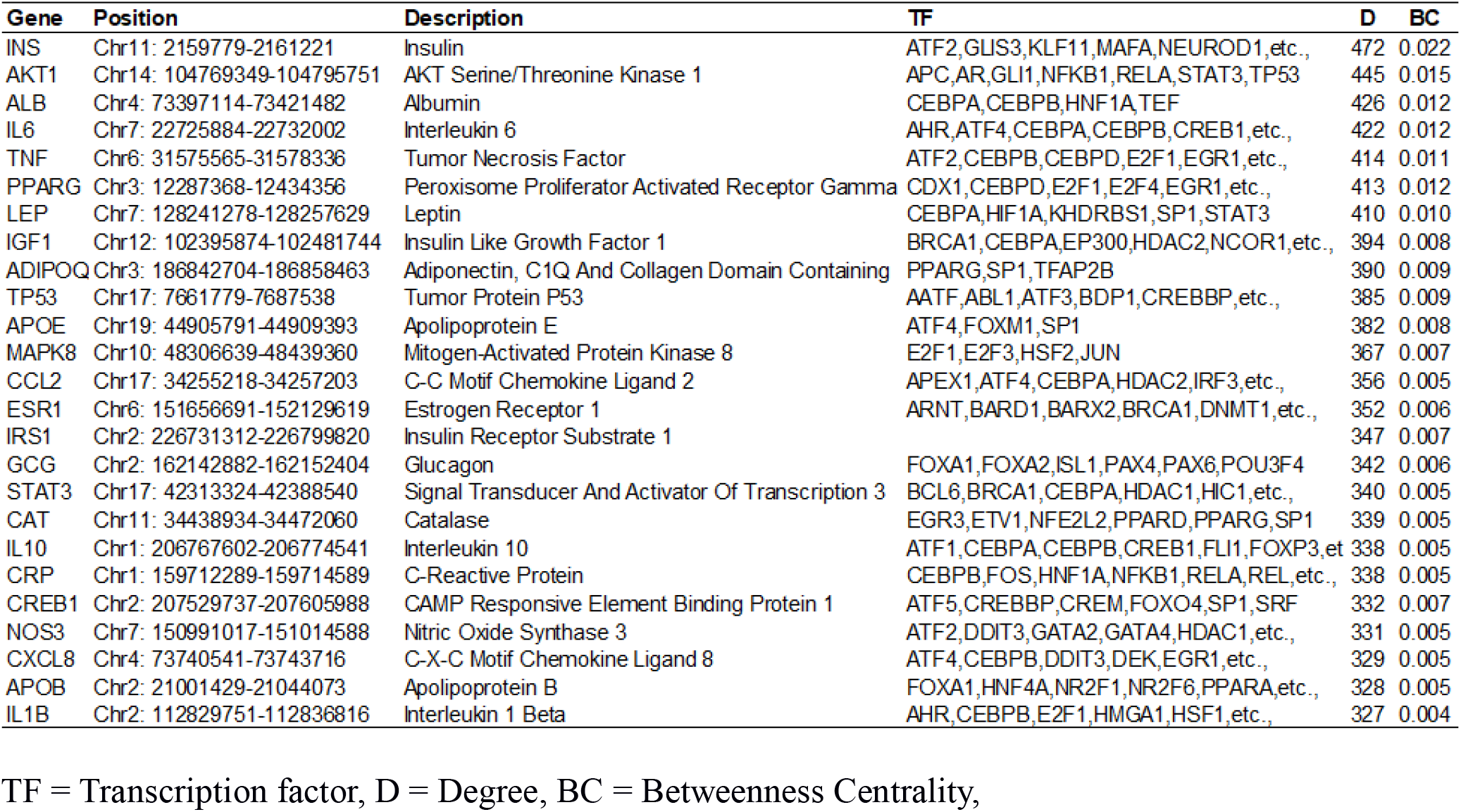
The 25 hub genes selected from the protein-protein interaction network.

**Figure 4:**
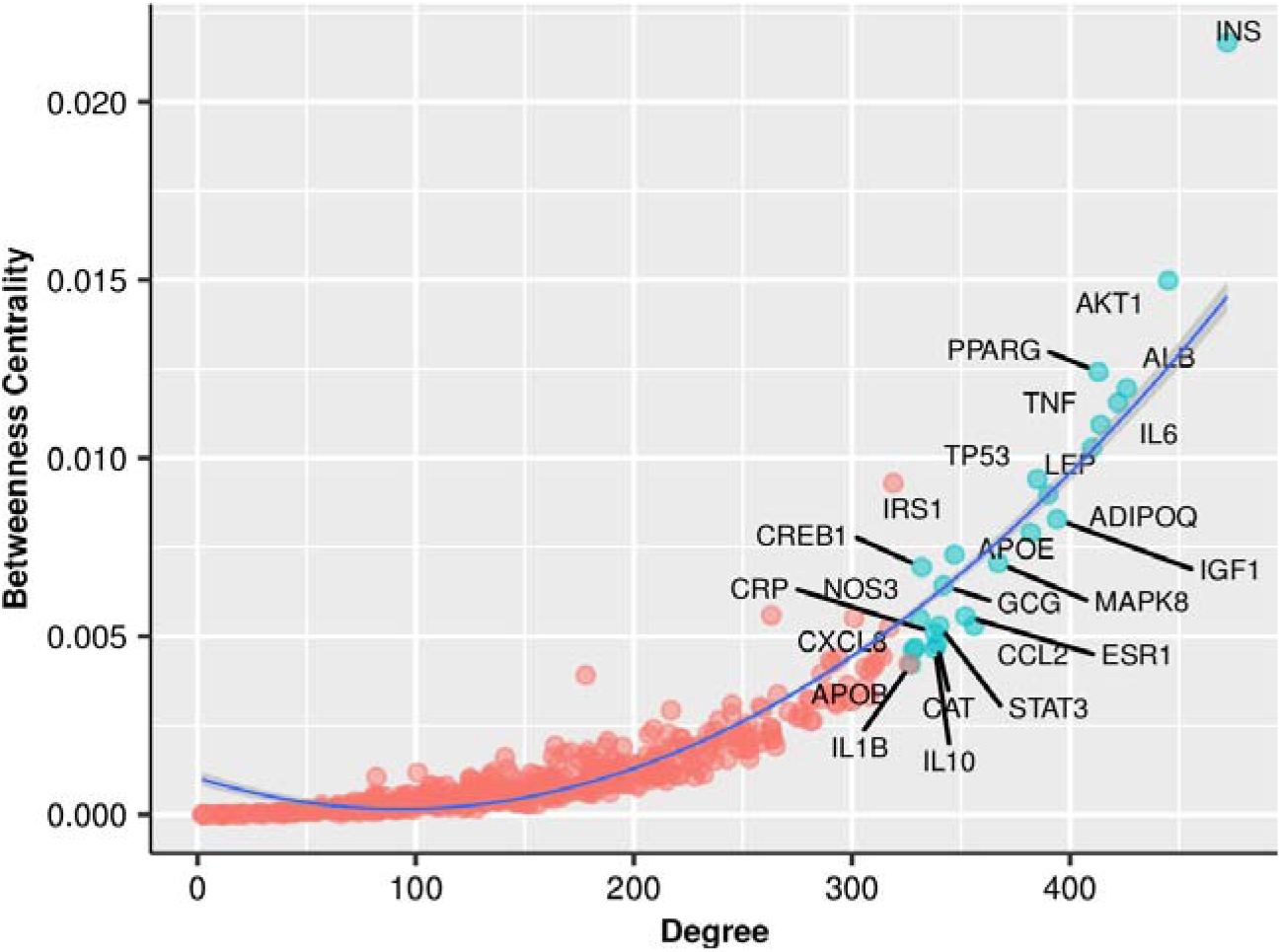
Relation between node degree versus betweenness centrality, hub genes were labeled and highlighted in blue color.

### Enrichment analysis of T2D associated genes

The Gene ontology (GO) enrichment analysis is a popular bioinformatics approach for determining if a set of candidate genes is enriched for certain biological processes (BP), molecular functions (MF), and cellular component (CC). The 273 genes involved in 81 enriched biological process GO terms passed the p-value and FDR cutoff. The top 25 highly enriched GO terms are shown as a dotplot in figure 5. The GO terms carbohydrate homeostasis (GO:0033500) and glucose homeostasis (GO:0042593) are the most significantly enriched biological processes with 34 number of genes involved in each process. Carbohydrate homeostasis refers to the balance between glucose production and utilization in the body to maintain normal blood sugar levels. In type 2 diabetes, there is a disruption in this balance, leading to imbalance in glucose homeostasis clinically referred as hyperglycemia (high blood sugar). The IRS2, GCK, LEP, IRS1, and ADIPOQ are the top five genes identified to have role in 58, 51, 50, 48, and 41 biological processes respectively. The IRS2 (insulin receptor substrate 2) is a protein involved in the insulin signaling pathway and its reduced function in type 2 diabetes can contribute to impaired glucose homeostasis. Understanding the mechanisms of insulin resistance and the role of proteins like IRS2 can help inform the development of new therapies for type 2 diabetes. A study conducted on IRS-2 deficient mice shows impaired both peripheral insulin signaling and pancreatic β-cell function which leads to progressive deterioration of glucose homeostasis [44]. In addition, some notable enriched biological processes linked to T2D associated genes include regulation of type B pancreatic cell apoptotic process, regulation of carbohydrate biosynthetic process, regulation of insulin secretion, regulation of hormone secretion, regulation of glucan biosynthetic process, regulation of glycogen biosynthetic process, regulation of peptide hormone secretion, hexose metabolic process, and monosaccharide metabolic process. The supplementary table 4 contains information related to all the enriched biological processes, molecular functions, and cellular components.

**Figure 5:**
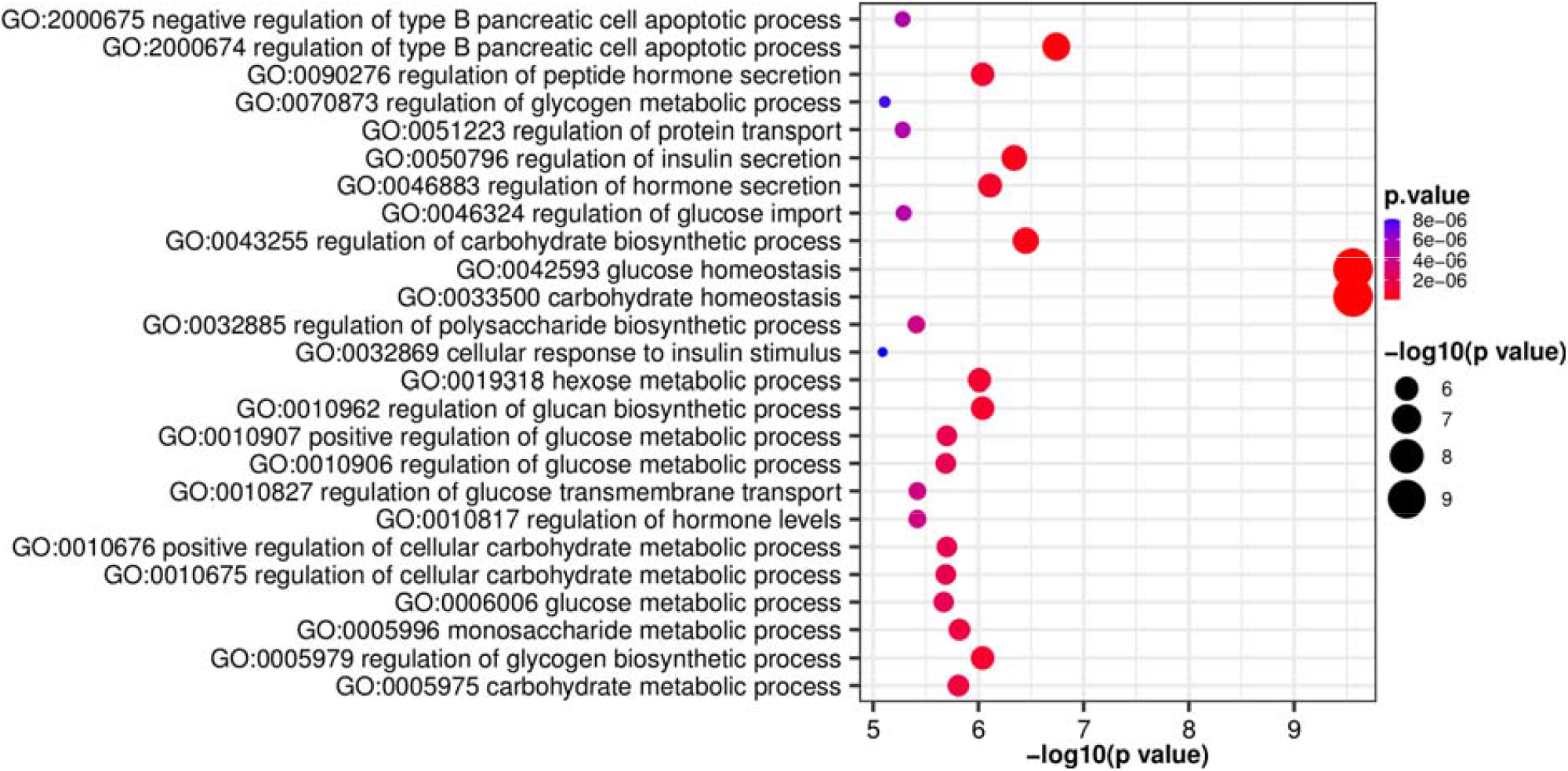
Gene Ontology (GO) biological process enrichment analyses of hub genes

To examine the involvement of hub genes in T2D further, we performed KEGG pathway enrichment analysis and found key enriched pathways with adjusted p-values <0.05. Insulin resistance, PI3K-Akt signaling pathway, AMPK signaling pathway, Neuroactive ligand-receptor interaction, Insulin signaling pathway, and MAPK signaling pathway are mainly enriched. The results for KEGG analysis are presented in figure 6 and corresponding genes in each enriched pathway are shown in Supplementary table 5.

**Figure 6:**
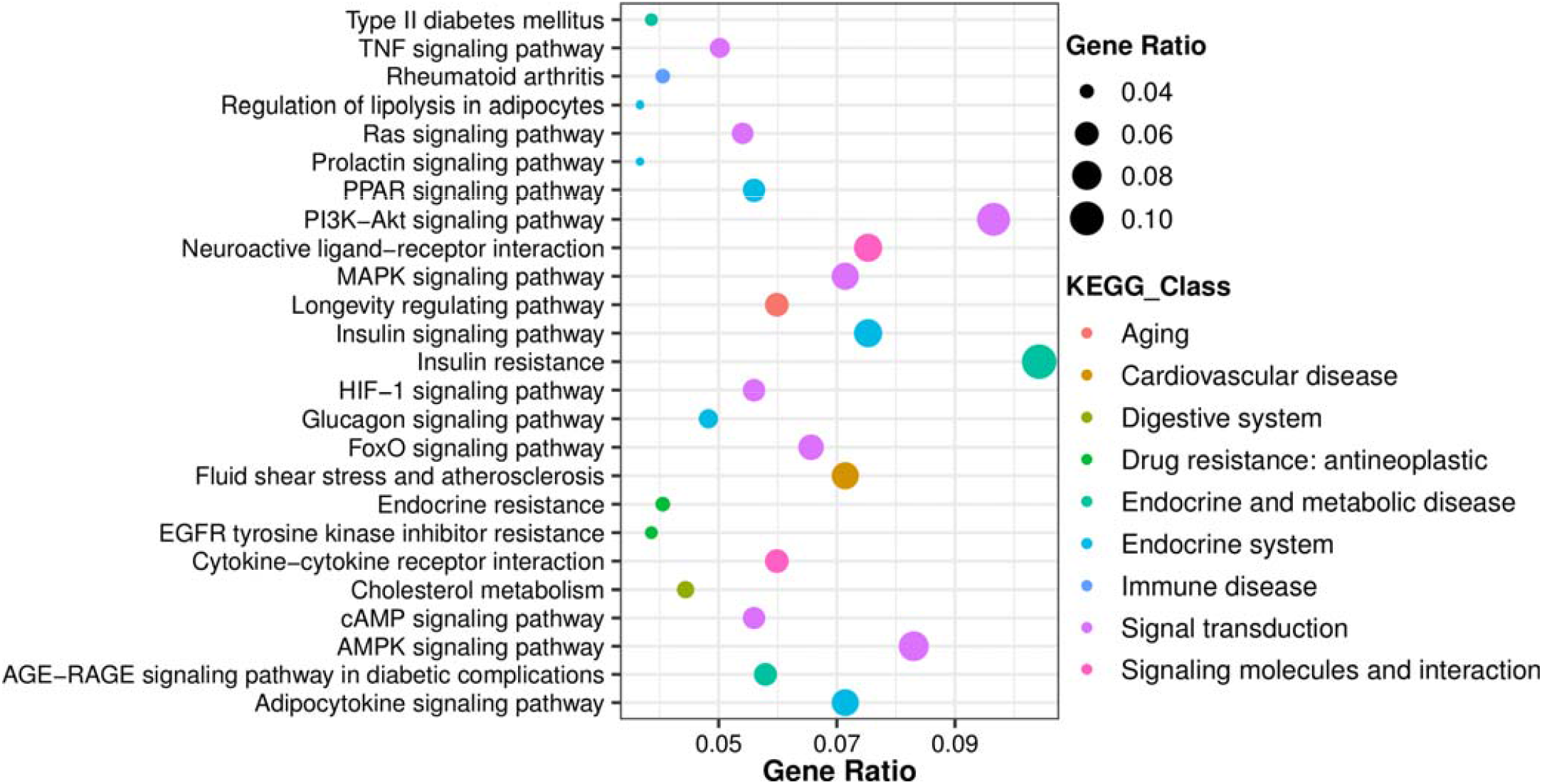
KEGG analysis of significantly enriched pathways of hub genes

### Prediction of miRNAs targeting hub genes

The regulatory relationships between the hub genes associated with T2D and their miRNAs were predicted using psRNAtarget (https://www.zhaolab.org/psRNATarget/) and established a interactive network using Cytoscape, which showed that the single gene may targeted by multiple miRNAs is shown in figure 7. All 25 hub genes were identified to have regulated by 54 different miRNAs belonging to 50 different families. The families miR-29, miR-125 and miR-1273 are having 3, 2, and 2 members in each family respectively. The remaining 47 families are possessing single members in each family. The hub gene IGF1 is targeted by 7 different types of miRNAs which are miR-1268b, miR-29a, miR-29b, miR-29c, miR-330-5p, miR-3934-5p, and miR-574-5p.

**Figure 7:**
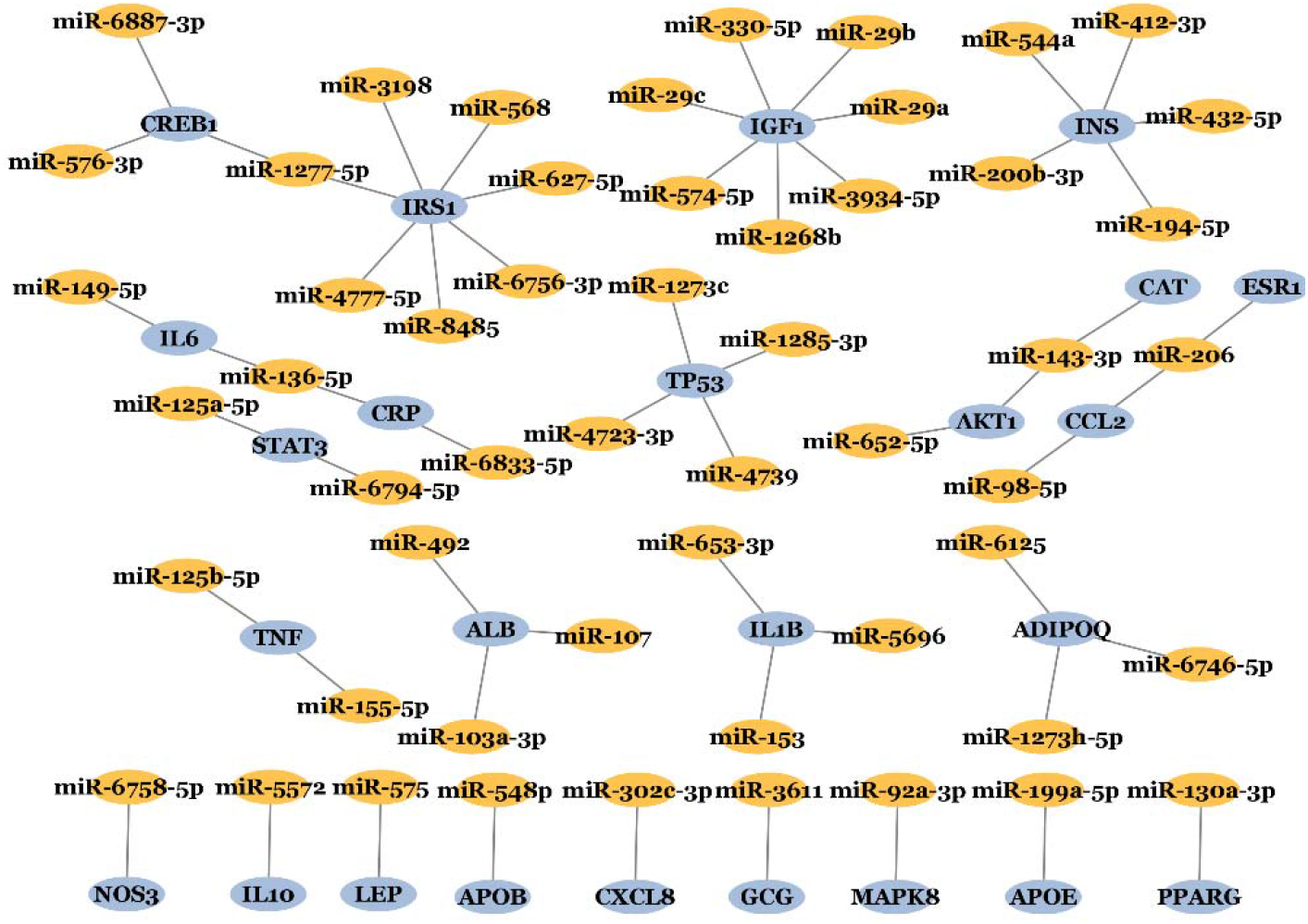
miRNAs targeting the hub genes were represented in the interaction network. Blue nodes represent hub genes in T2D, orange nodes denote miRNAs targeting hub genes.

The hsa-miR-1277-5p is targeting CREB1 and IRS1 genes. The IRS1 gene is targeted by 7 different miRNAs which include hsa-miR-568, hsa-miR-627-5p, hsa-miR-1277-5p, hsa-miR-3198, hsa-miR-4777-5p, hsa-miR-6756-3p, and hsa-miR-8485. Other miRNAs that have been implicated in type 2 diabetes include miR-126, miR-143, miR-375, miR-122, miR-103, and miR-107 [45] [46] [47]. These miRNAs have been shown to regulate various aspects of glucose and lipid metabolism, including insulin secretion, β-cell function, and hepatic lipid metabolism.

Three members of the miR29 family, miR-29a, miR-29b, and miR-29c, were identified as targeting IGF1, which is a growth factor that plays a critical role in cell growth, proliferation, and survival. Previous research found that miR-29a/b/c expression is higher in islets obtained from prediabetic non-obese diabetic (NOD) mice [48]. Increased miR-29a/b/c expression in islets and MIN6 cells can be triggered by proinflammatory cytokines or high glucose [49]. Most importantly, miR-29a/b/c exclusively target human and mouse MCT1, showing that miR-29a/b/c contribute to the β-cell-specific silencing of the MCT1 transporter and may provide a new therapeutic option for some manifestations of type 2 diabetes[50].

Our findings indicate that the miR136-5p has the potential to bind the 3’-UTR in the mRNAs of IL-6 (Interleukin-6) and CRP (C-reactive protein), are both markers of inflammation in the body [51]. They play roles in various physiological processes and are often associated with chronic conditions, including type 2 diabetes. The miR-136-5p appears to be involved in multiple aspects of diabetes, including insulin resistance, beta cell dysfunction, inflammation, and oxidative stress. However, our understanding of microRNA biology in diabetes is still expanding, and further study is needed to explain the precise processes and potential therapeutic implications of miRNAs in type 2 diabetes.

## Discussion

Pathological complications of type 2 diabetes remains undetermined and it has become a major public health concern worldwide [52]. Type 2 diabetes is a metabolic condition that affects how the body processes glucose, leading to high blood sugar levels. Over time, this can cause damage to various organs and systems in the body, leading to serious complications. The most common complications of type 2 diabetes include cardiovascular disease, kidney disease, nerve damage, eye damage and cognitive impairment. T2D not only causes physical and mental harm to those who suffer from it, but it also imposes huge financial burdens on individuals and nations. Since it is still a chronic disorder in modern medicine, thus identification of effective diagnostic and therapeutic markers is of utmost importance [53] [54]. In this study T2D associated genes and their annotations were largely retrieved and their corresponding protein-protein interaction network analysis was performed to find hub genes followed by their functional pathways were enriched. Ascertaining critical enriched pathways connected to T2D hub genes and their interactions will help better understand the illness and that may be prioritized for additional T2D clinical research.

Out of 490 associated genes selected for this study 25 hub genes were identified based on the top five percent of the genes with highest number of interaction with in the interaction network. The hub genes like IL6, IL1B, and IL10 are pro and anti-inflammatory cytokines dis-regulation of their levels are associated with increased risk of developing Type 2 Diabetes [55]. The finding in this study highlighted that regulation of peptide hormone secretion may play a crucial role in T2D, a recent study conducted on Leptin and Ghrelin documented that alterations in their ratios are associated with obesity and diabetes [56]. Leptin (LEP) and adiponectin (ADIPOQ) are peptide hormones that are produced by adipose tissue and play a role in the regulation of glucose and lipid metabolism. In individuals with type 2 diabetes, adiponectin levels may be decreased, leading to insulin resistance and impaired glucose metabolism [57]. Ghrelin is a peptide hormone that is produced by the stomach and stimulates appetite. In individuals with type 2 diabetes, ghrelin levels may be altered, leading to increased food intake and further exacerbating the metabolic abnormalities associated with the disease [58]. Some targets like IRS1, and IGF1 molecules are involved in the regulation of glucose metabolism and insulin signaling, and changes in their function or levels can contribute to the development and progression of type 2 diabetes [59] [60]. As clinical investigations progress toward therapies based on precision medicine, the information from this study may aid in the development of therapeutics based on genetic backgrounds.

The enrichment module determines whether pathways and GO keywords are statistically significant with the provided gene list by utilizing the over-representation analysis (ORA) method. The results of enrichment analysis of genes linked to type 2 diabetes revealed the central role played by biological pathways related to insulin resistance in the metabolic, pathological, etiological, and physiological aspects of the disease. Most of the enriched pathways had connection with the mechanism of insulin resitance, especially activation of PI3K-Akt signaling pathway, AMPK signaling pathway, Neuroactive ligand-receptor interaction and MAPK signaling pathway. The results of T2D-associated genes enrichment analysis revealed that carbohydrate homeostasis, and glucose homeostasis play a crucial role in the metabolism, disease progression, and physiology of type 2 diabetes.

In addition to identifying hub genes, we highlighted their target miRNAs in this work; about 54 miRNAs target all 25 hub genes. Differential miRNA expression contributes to the development and progression of type 2 diabetes by interfering with the aberrant regulation of their targets engaged in various stages of glucose homeostasis. We need a better understanding of the mechanisms underlying the abnormal production of these miRNAs in order to create viable therapeutics for type 2 diabetes. MicroRNAs’ significance in diabetes is becoming clearer, but more research is needed to identify their precise mechanisms and potential therapy implications in type 2 diabetes.

Overall, This study has identified multiple enriched pathways and their important hub genes using the knowledgebase workflow of Metascape. we have constructed a PPI network from T2D associated genes and have identified eight MCODE modules that may play important role in the pathophysiology of T2D. The identified 25 hub genes may serve as biomarkers of T2D and have the potential to be targets for therapeutic intervention. However, further studies are still needed to confirm our findings. These genes representing various enriched pathways of hub genes were observed to constitute a network of glucose metabolic process and insulin regulatory pathways. Even though it is hard to treat T2D patients based on these pathways and genes, future clinical studies can use the valuable information from this study to figure out the underling biomarkers in type 2 diabetes.

## Acknowledgments

The authors appreciate the support from Tumkur University in India.

## Author Contributions

This study was planned and conceived by PG. The complete bioinformatics analysis and manuscript drafting was done by PG. RS supervised this study and edited the manuscript. All authors contributed and approved the final manuscript.

